# Trends and Gaps on Philippine Scombrid Research: A bibliometric analysis

**DOI:** 10.1101/2021.06.30.450467

**Authors:** Jay-Ar L. Gadut, Custer C. Deocaris, Malona V. Alinsug

## Abstract

Philippine scombrids have been among the top priorities in fisheries research due primarily to their economic value worldwide. Assessment of the number of studies under general themes (diversity, ecology, taxonomy and systematics, diseases and parasites, and conservation) provides essential information to evaluate trends and gaps of research. This study presents a bibliometric evaluation of the temporal trends from 2000-2019 on Philippine scombrid research. Seven out of nine tuna species have SREA values equal to or higher than 1 with *Thunnus albacares* being the most researched tuna species. Twelve out of thirteen non-tuna species have SREA values less than one, thus indicating low research effort. It was apparent that there were significant differences in the number of studies in each thematic area except in ‘chemical analysis’ and ‘diseases and parasites’ between scombrid groups where there was low research effort observed. The research points at the uneven research distributions between scombrid groups in each thematic area. As locally published research are significantly behind foreign publications in terms of citation index, international collaborations by Filipino researchers have shown an increase research impact. Our study hopes to influence the local and international R&D agenda on Philippine scombrids and promote solidarity among nations towards its conservation and management.

## INTRODUCTION

The Philippines, the second-largest archipelago in the world, is composed of 7,641 islands. As an archipelago, the country is surrounded by major and minor seas, which comprise frequent spawning areas in coral and pelagic regions. The country’s archipelagic formation combined with a wide sea area contributes to its vast fisheries resource (Catibog-Sinha, 2010). The geographic feature of the country led most Filipinos to have a high dependence on marine resources for livelihood and nutrition. In addition to climatic conditions and archipelagic formation, the phenomenal forcing of monsoonal winds causes upwelling, which significantly contributes to the high productivity of every local fishery (Licuanan et al., 2019). Thus, with these favorable conditions, the Philippines ranked ninth globally in fisheries production in 2017 (BFAR, 2019).

Studies have recorded the existence of 22 scombrid species in the Philippine seas. Such vast resources and biodiversity underlay why this family is considered as one of the most economically important marine species in the country (http://www.fishbase.org). The scombrid resources of the Philippines have high economic standing not only because they generate export revenues but also provide a major protein source for local and foreign consumption (SEAFDEC, 2017). The country was reported to export more than 170 million metric tons of tuna globally, which made the Philippine the top 2 global tuna producer in 2018 (BFAR, 2019). Furthermore, in terms of non-tuna scombrid production, the country was also the 2^nd^ top mackerel producer in Southeast Asia in 2014 (SEAFDEC, 2017).

Unfortunately, the world’s stock status of scombrids has seen significant reduction over the past fifty years (Annala & Eayrs, 2010). Yang & Pomeroy (2017) reported that the trend was even worse in the Philippine seas due to some oblivious factors. The rate of deterioration was influenced by critical issues that include overfishing, poor management, small and large fisheries conflicts, habitat degradation, lack of research and information, and inadequate technical and human capabilities. Despite this trend, there has been no comprehensive assessments focused on this family, thereby contributing to incomplete and fragmented knowledge of how these economically important fishes can be sustainably managed or conserved. Consequently, there is an apparent need to assess the trends to calibrate R&D effort and strategies at local and international areas (Mallari et al., 2015). Thus, the analysis of existing information on scombrid in Philippine waters is critical first step towards the recrafting of a marine fisheries R&D agenda (Juan-Jordá et al., 2013). This study presents a bibliometric analysis of literature on scombrids from Philippine waters from 2000-2019. In particular, the analysis identities the patterns and trends in thematic areas of research, research effort allocation on specific scombrid species, publication influence, authorship, and research collaboration. Such information might provide fresh insights into the conglomeration of scientific behavior and dynamics in research priorities that could benefit both the scientific community and policy developers.

## METHODS

### Data Collection

The publications were collected from Google Scholar (https://scholar.google.com), PubMed (https://www.ncbi.nlm.nih.gov/pubmed), and ScienceDirect (https://www.sciencedirect.com) The search formula used was “Philippine” AND (“Scombrids” OR “Scombridae” OR the different scientific names). The scientific names used in the search are: *Acanthocybium solandri, Grammatorcynus bilineatus, Gymnosarda unicolor, Rastrelliger brachysoma, Rastrelliger faughni, Rastrelliger kanagurta, Sarda orientalis, Scomber australasicus, Scomber japonicas, Scomberomorus commerson, Scomberomorus guttatus, Scomberomorus queenslandicus, Scomberomorus semifasciatus, Auxis rochei, Auxis thazard, Euthynnus affinis, Katsuwonus pelamis, Thunnus alalunga, Thunnus albacares, Thunnus obesus, Thunnus orientalis*, and *Thunnus tonggol*. A total of 158 articles were collected from Google Scholar (N=84), PubMed (N=13), and ScienceDirect (N=27) indexed from January 2000 to December 2019. The results were filtered for overlaps or duplications, resulting in 124 articles. All filtered publications from search results were categorized according to species groups, publication years, and authors’ origin information. To assess scombrids’ research allocation, we collated the literature according to main thematic areas that include (1) Diversity, (2) Ecology, (3) Systematics, (4) Chemistry, (5) Diseases, and (6) Conservation. This kind of approach is an effective measure of allocation of national, global, or regional conservation efforts (Meyer et al., 2015). The main thematic areas were further divided into secondary categories as described in Table 1 to refine and differentiate all studies into more specific areas. The profiles of international cooperation were also analyzed based on the affiliations of the authors. Data on the funding source were derived from the “Acknowledgement” section and funding source information provided by the article, whenever available.

**Table I.**
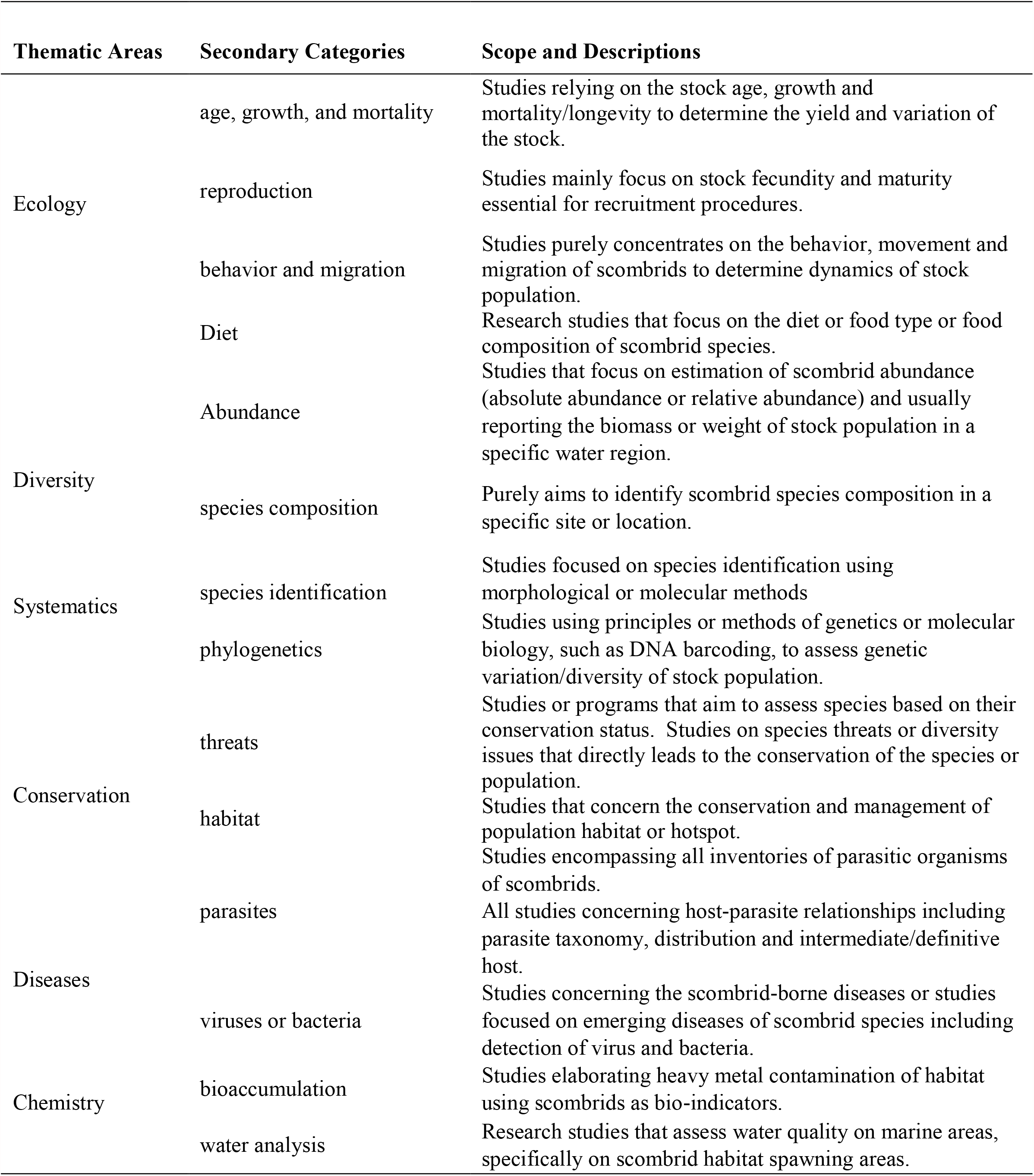
Thematic Areas and secondary categories of research reviewed based on categories from Tanalgo and Hughes (2018)

### Data Analysis

Data obtained were exported to a database (MS Excel, Microsoft Corp., USA) where pertinent descriptive statistics were used. All statistical analyses were performed using Statistica v 10 (StatSoft Inc., 2011) and PAST ver. 3.18 (updated version 2018) respectively. Significance was set at *p*=0.05.

*Species-Research Effort Allocation (SREA)*. We adopted the species-research effort allocation (SREA) metric to quantify research efforts among species temporally. SREA is a simplified metric that allows identifying species or taxonomic groups that have received adequate attention in a certain period. According to Tanalgo et al. (2018), ideally, the SREA metric is effective in a review covering a more extended period (e.g., more than ten years).

SREA is expressed using the equation: SREA (x) = R□/y

> where: x = species or scombrid group; R□ = species records (number of times species was recorded from all publications/reports); and y□ = the number of years covered by the review or assessment

Species-Research Effort Allocation (SREA) value equal to 1.00 indicates an average effort per year relative to all species. In contrast, SREA >1.00 indicates a higher effort was given to the species, and <1.00 indicates a lower research effort. SREA values was also used to test the difference of the number of studies among scombrid groups (Tuna and Non-Tuna) in broad and thematic areas (Diversity, Biology, Taxonomy and Systematics, Diseases and Parasites, Chemical Analysis and Conservation) using Mann-Whitney U-test (Fowler et al. 1998).

### Equitability Index

The Shannon evenness index (J’) was calculated to assess the equitability of research between scombrid groups (tuna and non-tuna) on different thematic areas. In this study, J’ was computed as: J’ = SDI / max (SDI) = - ∑ (Pi * In (Pi)) / ln(m)

> ; where the value of J’ is constrained between 0 and 1, which is interpreted as values approaching 1 indicates an equal proportion of research allocated.

Pearson’s Chi-squared test of independence (χ2) was used to test the difference in the proportion of studies in broad thematic areas and between the scombrid groups.

### Citation Index

Citation index for each paper and author were obtained through Google Scholar as it yields more robust results. Average citations per year for each thematic area and the scombrid group were calculated based on the total citations per year multiplied by the total number of papers.

## RESULTS AND DISCUSSION

### Philippine scombrid research based on the thematic areas

The study has searched and reviewed 124 online publications for scombrid research from 2000 to 2019 based on six target thematic areas: Diversity, Ecology, Taxonomy and Systematics, Chemical Analysis, Diseases and Parasites, and Conservation. **Table 1** presents the various thematic areas and secondary categories of research reviewed with some modifications (Tanalgo & Hughes, 2018). All six thematic areas provide crucial information for future conservation and management strategies. There was an average of 6.2 publications on scombrid research annually from 2000-2019.

The 124 publications consisted of minimal differences in records for Diversity, Ecology, and Conservation themes. Thus, these publications contributed overlapping themes on each publication for the findings that one paper may have covered several themes. It was found out that ecology-related publications (n=59, 47.58%) were the most dominant followed by themes on diversity (n=55, 44.35%) and conservation (n=51, 41.13%). However, every thematic area was subdivided into several specific secondary categories to evaluate the more specific focus of each publication based on the content of the paper.

While ecological studies (n=59) were subdivided into four categories, we noted that the secondary categories within this thematic area have frequent overlaps. Nine publications were on scombrid’s ‘diet,’ 39 publications discussed ‘age, growth, mortality or longevity,’ 17 publications were on ‘reproductive biology,’ and 27 publications assessed ‘behavior, migration, and movements.’ The ‘diet’ studies mainly evaluated the tuna scombrid group’s feeding type in this thematic area. For instance, Baeck et al. (2014) reported that *Auxis rochei* in Ilo-ilo, Philippines, is an epipelagic offshore predator feeding on whatever abundant resources are available in the environment, have a narrow food niche, and is a specialized feeder with fish as their dominant prey. Ecological studies were overshadowed by the sub-theme ‘age, growth, mortality and longevity,’ and publications from this category were focused on assessments primarily conducted for fishery purposes for stock population evaluations of scombrids. However, this category emphasized fish stocks estimation before any actual collection. Also, studies under this secondary category were technically and mostly aided by the length based on the MULTIFAN method. The MULTIFAN is a likelihood-based method for estimating growth parameters and age composition from multiple length frequency data sets. This method has been successfully applied to estimate the age and growth of various marine species such as Bluefin tuna, Albacore tuna, and Skipjack tuna (Su et al., 2003). The ‘reproductive biology’ studies consisted mainly and more focused on the tuna group. The leading objective in this category was to define the spatiotemporal and size-related patterns in reproductive parameters of scombrids (Itano, 2000). The ‘behavior, migration and movement’ studies were also more focused on the tuna group and predominantly conducted on the Philippine Pacific region, South Taiwan waters, and the West Philippine Sea. Most observations from this category provide essential insights into habitat use, especially in tuna (Macusi et al., 2015) around Philippine waters.

Diversity studies (n=55) were split into two categories. Publications on diversity assessments comprised 47 publications that discussed ‘abundance’ and 34 publications for ‘species composition.’ This trend shows that diversity studies on Philippine scombrids were dominated by abundance assessments and reports on payao or some fish aggregating device in the Visayas waters. In these studies, purse seines and ring nets aided in harvesting large-sized tunas, small tunas, and small pelagic species (Babaran & Ishizaki, 2010). In addition, Babaran & Ishizaki (2010) considered that these kinds of gear differ in terms of the scale of operations, level of mechanization, and mode of the fishing operation, but essentially capture the same species of a similar size range. However, most ‘species composition’ information was based on the catch compositions on specific regions of Visayan waters performed by fisheries departments. Species composition of scombrids is also affected by seasonal variation (Baleta & Baleta, 2016).

Conservation studies (n=51) were also subdivided into two categories. This theme was composed of studies to assess and strengthen the conservation and management efforts of scombrids. Under this thematic area, overlaps of secondary categories on publications were recorded. It was assessed that 46 publications focused on understanding the status of ‘species and threats’ and 27 publications that reviewed ‘habitats.’ Furthermore, all the publications on conservation and management of fisheries overlapped with most thematic areas. The conservation and management topics are mainly based on fisheries departments’ information. Most data based on ‘species and threats’ was due to the overfishing techniques that decline the number of the stock population, including the larvae species. Also, based on ‘habitat,’ one central point of all studies under this category was the illegal fishing methods that aggravated the degradation and destruction of most spawning areas essential for scombrid reproduction.

Taxonomy and Systematics studies (n=29) were partitioned into two categories containing 17 papers on ‘species identification’ and 19 concerning ‘phylogenetic’ analyses. The ‘species identification’ category under this thematic area has mainly utilized mitochondrial DNA for barcoding for scombrid species identification purposes and verification of new scombrid species existence within Philippine waters primarily performed on tuna scombrid group. Additionally, species identification in the non-tuna scombrid group utilized mitochondrial DNA through PCR-RFLP analysis (Muto et al., 2017).

Chemical Analysis (n=7) was subdivided into two categories. All these publications covered topics for scombrid ‘bioaccumulation.’ Moreover, four publications included topics assessing ‘water analysis’ for scombrid spawning marine areas. However, this thematic area containing both the categories was the least studied thematic area for scombrid research throughout 20 years. It only had seven publications - a minor proportion (5.65%) than other thematic areas. ‘Bioaccumulation’ studies were found to be conducted only on tuna. Without exception of scombrids, heavy metal contaminants are a persistent problem by all fish species (Dacera et al., 2018). Also, studies covering ‘water analysis’ were based on bottom sediments to track the possible sources of contaminants (Dacera et al., 2018). These studies were all conducted on local waters within Mindanao and Visayas.

When it comes to other thematic areas, disease and parasites (n=10) were also subdivided into two categories, namely ‘parasitic association’ and ‘viral and bacterial associations.’ Eight parasite-relevant publications highlighted acanthocephalan parasites association on scombrids. Parasites from the Phylum Acanthocephala encompass worms with various recorded sizes and many intermediate, paratenic, and final hosts. There were only a handful of papers in this category because Acanthocephalan parasites are rare in Philippine waters (Briones et al., 2015). Unfortunately, studies on ‘viral and bacterial associations’ are very few. Therefore, this field must be given more effort to fill the research gaps. Figure 1 summarizes the number of studies in all specific secondary categories under six thematic areas and the profiles of cross-cutting research.

**Figure 1.**
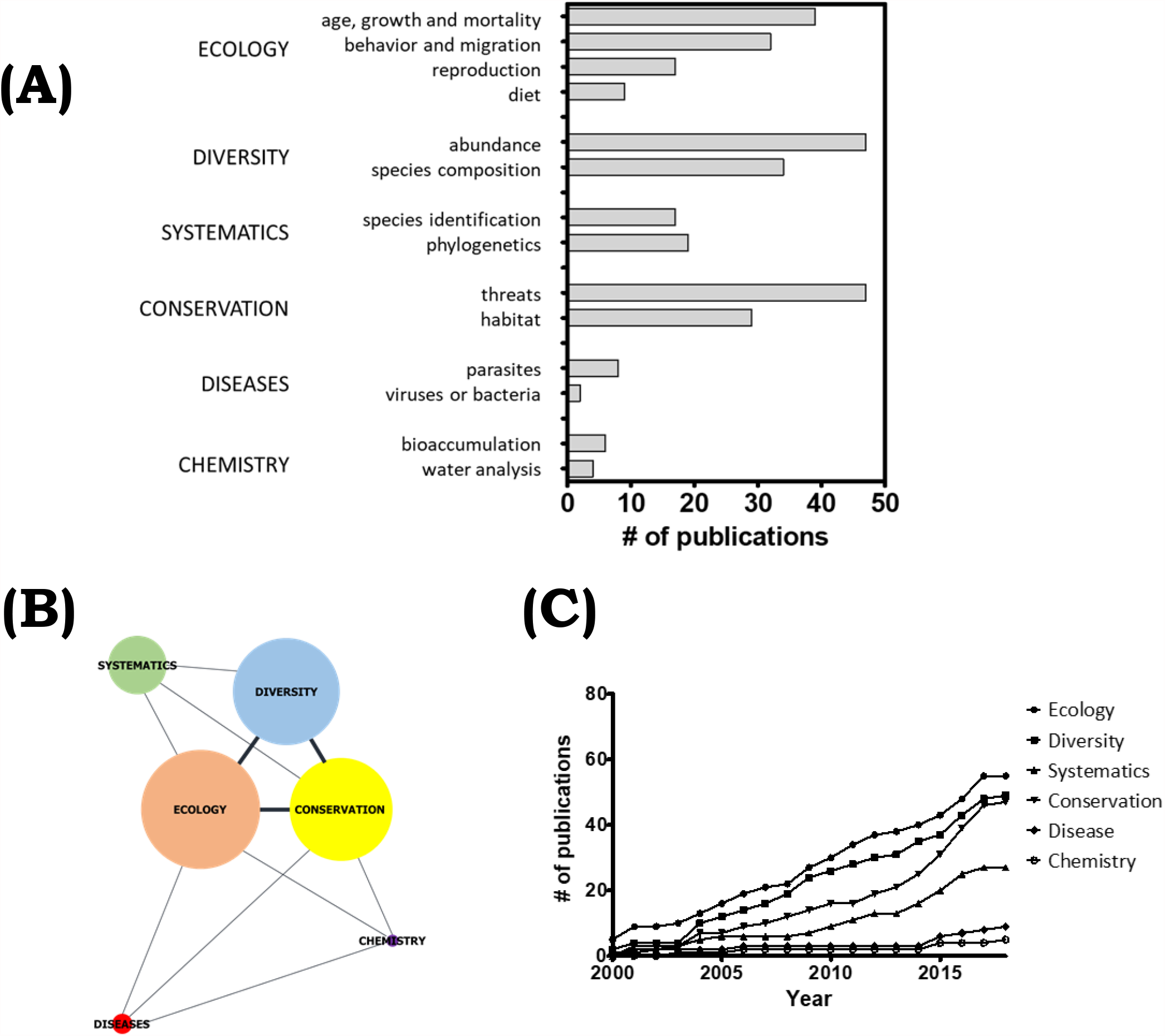
An assessment of published papers on Philippine Scombrids. **(A)** The number of studies published in six thematic areas with corresponding secondary categories, **(B)** the profile of studies spanning multiple thematic areas, and **(C)** the number of papers published from 2000-2019.

### Temporal variation of research themes

There was an average of 6.2 publications per year on scombrid research from broad thematic areas. The highest numbers of publications covering all six thematic areas were in 2017 and 2016. Ecological-relevant reports (n=59, 47.58%) had the highest number among all thematic areas with an average of 2.95 publications per year except in 2002 and 2018. Diversity-related publications (n=55, 44.35%) gave an average output rate of 2.75 papers per year, while conservation-related reports (n=51, 41.13%) had an average annual publication rate of 2.55.

These three main thematic areas had an average annual publication of more than two compared to the other thematic areas. For example, papers on chemical analysis (n=7, 5.65%) only appeared throughout the 20 years had only an average annual publication rate of 0.35). Figure 1C demonstrates the timeline of increasing publications from 2000-2019 based on specific thematic areas.

### Species Records and SREA

There are 428 scombrid species records, though the non-tuna group is more biodiverse. All searched publications were assessed to determine the number of species records (or species occurrences from the publication sets) from the two scombrid groups. The results revealed that the number of species records of the tuna group (n=301) was more numerous than those in the non-tuna group (n=127). Since Species-Research Effort Allocation (SREA) values of most species under the tuna group were higher than 1, it was apparent that research activities were more focused on tuna species. Unfortunately, species from the non-tuna group were evaluated to have lower values of SREA. Figure 2 illustrates SREA values of the scombrid species.

**Figure 2.**
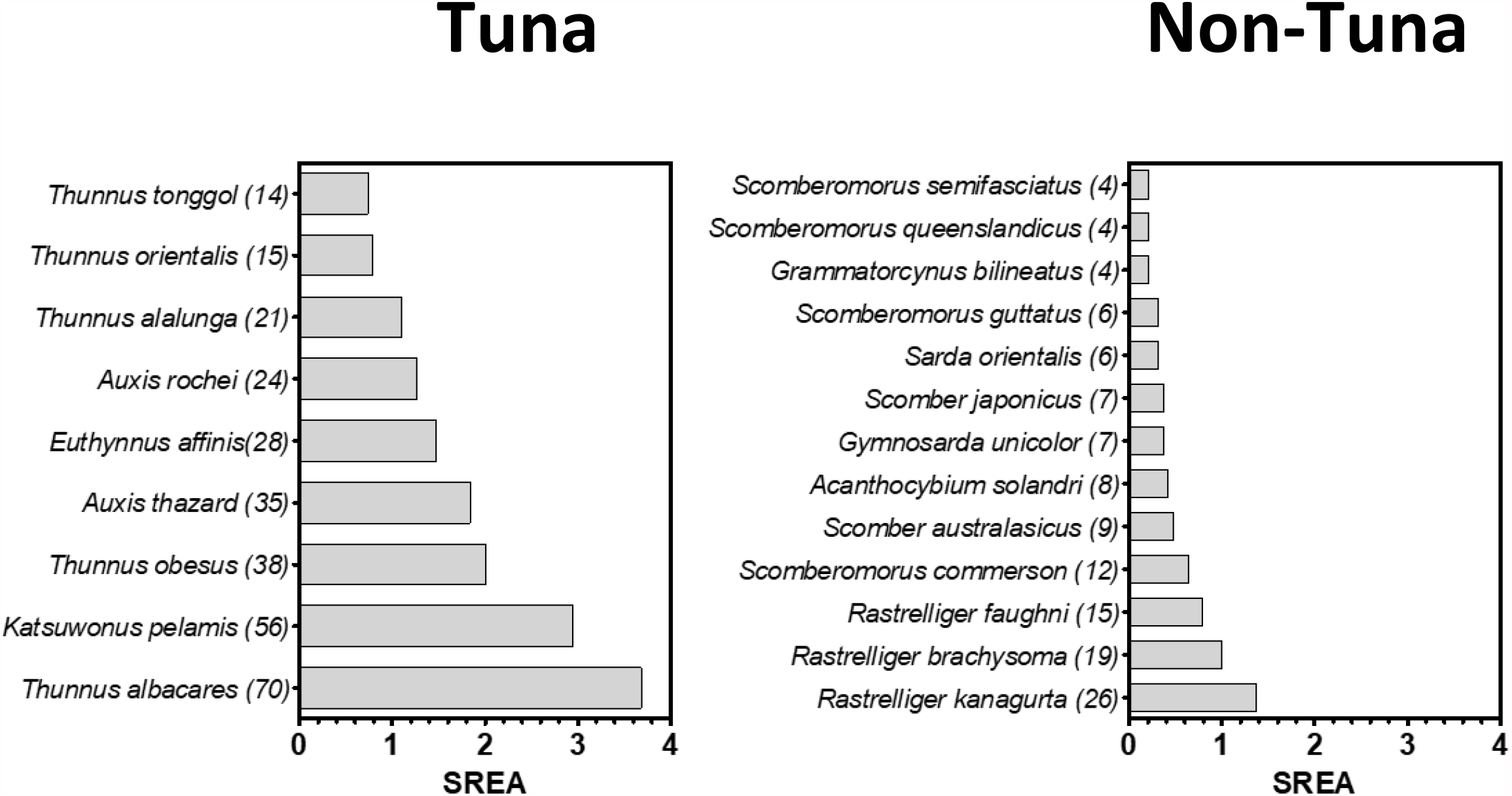
Species-Research Effort Allocation (SREA) of Philippine scombrid species. **(A)** Tuna and **(B)** non-tuna species. The majority of the non-tuna species are below the average (dash line) SREA suggesting a lack in research allocation or activity.

As expected, species records under the tuna group (n=301) were most frequent in diversity-related studies (n=152). This theme was followed by ecological studies (n=103), conservation (n=72), taxonomy and systematics (n=25), diseases and parasites (n=11) and chemical analysis (n=8). Among the nine tuna species, seven have SREA values of more than 1, with most research on *Thunnus albacares*, commonly known as Yellowfin tuna (SREA=3.68). *Katsuwonus pelamis* (Skipjack tuna; SREA= 2.95), *Thunnus obesus* (Big-eye tuna; SREA= 2), *Auxis thazard* (Frigate tuna; SREA= 1.84), *Euthynnus affinis* (Kawakawa; (SREA= 1.47), and *Auxis rochei* (Bullet Tuna; SREA= 1.26) all received high research allocation. These tuna species are also considered one of the most traded seafood commodities globally. Pecoraro et al. (2018) also reviewed that despite its great biological and economic importance, there was increased interest in classical genetic and tagging studies of Yellowfin tuna population structure at local and global oceanic scales. However, two tuna species, *Thunnus orientalis* (Pacific bluefin tuna) and *Thunnus tonggol* (Longtail tuna) were found to have SREA values lower than 1. Notably, *Thunnus orientalis* is considered a commercially important species. Still, it is listed as ‘vulnerable’ on the International Union for Conservation of Nature (IUCN) due to overfishing (Sarmiento et al., 2016).

Compared with the tuna group, there were fewer species records (n=127) with the non-tuna scombrids. Most frequent species citations were seen in diversity studies (n=59), followed by ecology (n=24) and conservation (n=16). Among 13 species of the non-tuna scombrid group, only *Rastrelliger kanagurta* (Indian mackerel) had an SREA value higher than 1 (SREA=1.37). The Indian mackerel has a worldwide distribution that forms an important part of the fishery industry in Southeast Asia. Pedrosa-Gerasmio et al. (2015) underscore its species ubiquity may have contributed to the immense attention on ecological assessments of this species. The rest of the 12 species were found to have SREA values lower than 1, signifying the need for more research attention.

Based on our analysis, there were more publications on diversity (U=14.5, *p=*0.002), ecology (U=7, *p=*0.0003), conservation (U=24.5, *p=*0.01), and taxonomy and systematics (U=23.5, *p=*0.01) in favor of the tuna compared to the non-tuna scombrids. Since papers on chemical analysis containing any of the species were nominal in the past 20 years, no significant difference was seen between the scombrid groups (U=39.5, *p=*0.11). Species allocation was also ‘balanced’ in diseases and parasite studies between the scombrid groups (U=54.5, *p=*0.41). Combining all SREA values of each species from broad thematic areas, it was clear that the scombrid tuna group received significantly more research attention than the non-tuna group (U=6.5, *p=*0.0003) in the broad thematic areas. Moreover, the species records between tuna and non-tuna scombrids from broad thematic areas significantly differ (χ2=25.87, *p=*0.001), indicating an imbalanced proportion of studies. Finally, there was uneven research distribution specifically on individual thematic areas regarding the evenness or equitability of research efforts. “Diversity (J’=0.84), Ecology (J’=0.63), Taxonomy and Systematics (J’=0.82), Chemical Analysis (J’=0.59), Diseases and Parasites (J’=0.0.81) and Conservation (J’=0.76) studies all have uneven research efforts distribution between scombrid groups.

### Researcher Profiles and Collaborations

Based on the analysis conducted, 451 authors produced 124 publications for scombrid research within Philippine waters from 2000-2019. A closer look at the authors’ affiliations, 48.4% of the papers were from foreign institutions, while 29.8% were from Philippine-based researchers. Less than a quarter (21.8%) of the studies on scombrids in Philippine waters were a collaboration by local and foreign researchers.

The collaborative studies between Filipino and foreign research were mainly through the Southeast Asian Fisheries Development Center (SEAFDEC). Interestingly, the joint research initiatives were more focused on conservation issues. Figure 3 summarizes the research engagements of local, foreign, and collaboration between local and foreign researchers on scombrids in Philippine seas based on thematic areas and scombrid groups.

**Figure 3.**
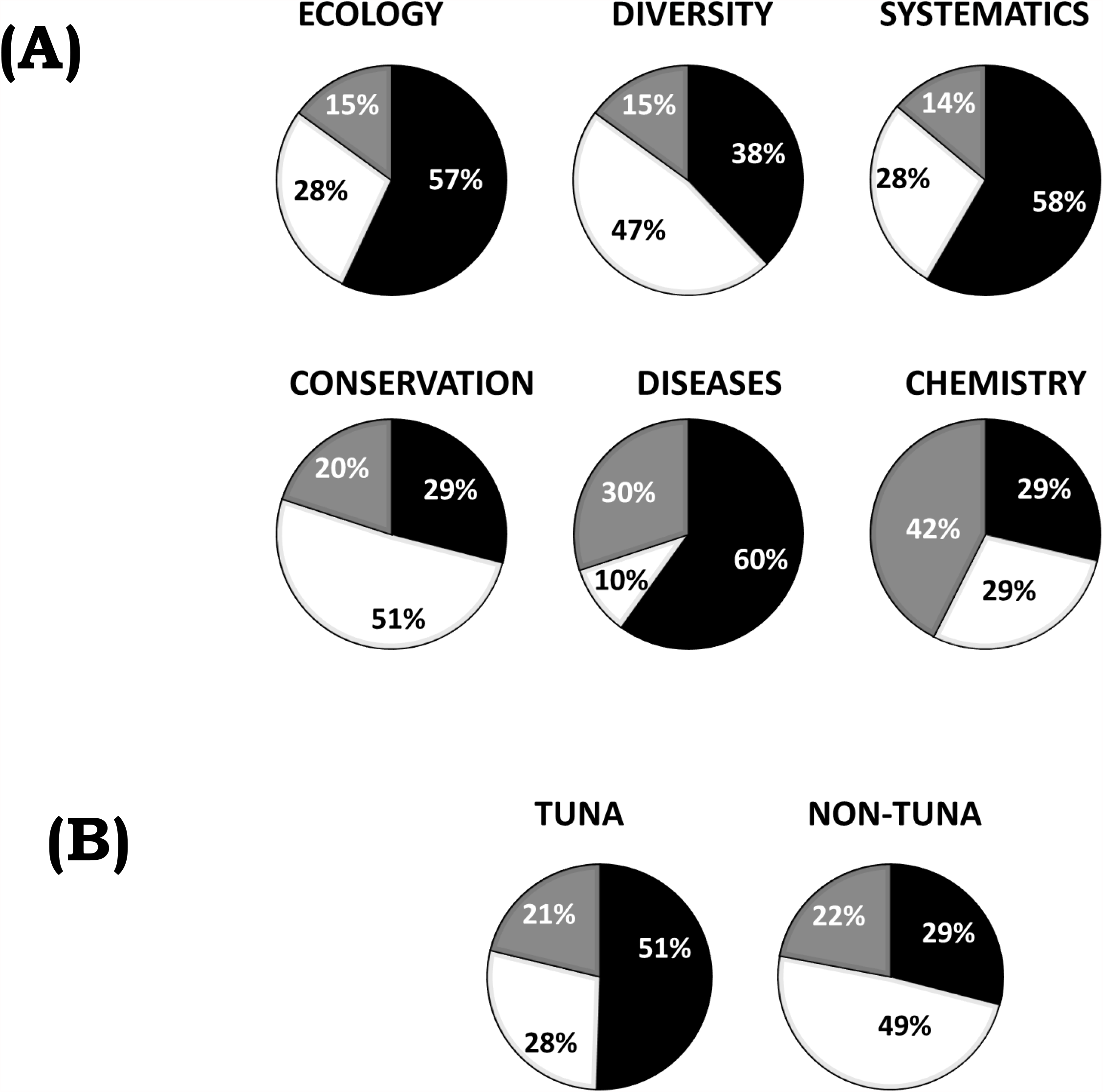
Pie charts of research engagements on scombrids in Philippine seas. The charts show allocation of research according to (A) thematic areas and (B) scombrid group. Studies were conducted by foreign groups (black), local researchers (white), and collaboration bet local and foreign researchers (grey).

Finally, we assessed the citation index of a papers. The citation index measures the importance of the index paper within the scientific community. It reflects the impact of a published material where a work of great significance is likely to be cited more often. The total number of citations for a published paper refers to the number of scientific articles citing it in their bibliography. As shown in Table 2, documents from foreign-based researchers generated significantly (*p=*0.001) higher citation indices in all thematic areas and across the scombrid groups, even if there were instances where local researchers produced more papers. For example, while there were more papers published by Filipino researchers in the field of diversity, foreign authors garnered a much higher citation index (1.15) compared to the locals (0.35). On the other hand, we observed that papers co-authored by both foreign and local researchers generated higher citation indices compared to those solely by Filipino researchers.

**Table II.** Average citations per year (N, number of papers) of research engagement on scombrids in Philippine seas.

|  |  | Local<br>(Philippines) | Foreign<br>(single or multi-country) | Philippine International<br>Collaboration |
| --- | --- | --- | --- | --- |
| **Thematic Area** |  |  |  |  |
|  | Ecology | 0.19 (16) | 1.79 (33) | 1.45 (9) |
|  | Diversity | 0.32 (26) | 1.15 (21) | 0.13 (8) |
|  | Systematics | 0.58 (8) | 1.57 (17) | 0.47 (4) |
|  | Conservation | 0.47 (26) | 0.83 (15) | 0.53 (10) |
|  | Diseases | 0.02 (1) | 2.88 (6) | 0.49 (3) |
|  | Chemistry | 0.13 (2) | 1.03 (2) | 1.29 (3) |
|  |  | 79 | 94 | 37 |
| **Scombrid group** |  |  |  |  |
|  | Tuna | 0.25 (31) | 1.53 (55) | 0.41 (23) |
|  | Non-Tuna | 0.35 (20) | 0.97 (12) | 0.47 (9) |
|  |  | 51 | 67 | 32 |

We should bear in mind that aside from the type of research engagement, there are several factors that influence the citation index of a paper. One is online availability or open access. Second is the data source of the documents. Citation indices acquired from Google Scholar are likely to be higher than those obtained from other indexing services like Scopus and Researchgate (Sarkar & Seshadri, 2015). The third is the total volume of scientific publications in that area. A ‘hot field’ is more likely to gain more citations. And lastly, senior researchers are likely to be cited more often because of their reputation and established research network within the scientific field. Table 3 summarizes the top 10 authors with the most publications, and Table 4 summarizes the top 50 most cited articles in scombrid research from 2000-2019. The ranking was based on the annual citations per year. Note, this may provide a birds-eye-view of the top authors and papers and may not be equated in terms of scientific impact as there may be less propensity for an article to be cited or noticed if published recently.

**Table III.** Top 10 authors with the most number of publications from 2000-2019.

| Author | # of Papers | Affiliation |
| --- | --- | --- |
| John Hampton | 10 | Oceanic Fisheries Program, Secretariat of the Pacific Community, Noumea, New Caledonia |
| Ricardo P. Babaran | 8 | College of Fisheries and Ocean Sciences, University of the Philippines Visayas, Ilo-ilo, Philippines |
| Mudjekeewis D. Santos | 7 | Genetic Fingerprinting Laboratory, Research and Ecological Assessment Division, National Fisheries Research and Development Institute, Quezon City, Metro Manila, Philippines |
| Adam Langley | 6 | Oceanic Fisheries Program, Secretariat of the Pacific Community, Noumea, New Caledonia |
| Rafael V. Ramiscal | 4 | Bureau of Fisheries and Aquatic Resources 860 Quezon Avenue, Arcadia Bldg., Quezon city, Philippines 1103 |
| Noel C. Barut | 3 | Marine Fisheries Research and Development Division, National Fisheries Research and Development Institute, 940 Quezon Ave., Quezon City 1103 Philippines |
| Angel B. Encarnacion | 3 | Department of Agriculture-Bureau of Fisheries and Aquatic Resources Regional Office No. 2 (Regional Government Complex, Carig, Tuguegarao City 3500 Cagayan, Philippines) |
| Chikayuki Endo | 3 | Faculty of Agriculture, Kinki University, 3327-204 Naka-machi, 631-8505 Nara, Japan |
| David G. Itano | 3 | National Marine Fisheries Service, Pacific Islands Region, 1601 Kapiolani Boulevard, Suite 1110, Honolulu, HI 96814, U.S.A |
| Pierre Kleiber | 3 | Pacific Islands Fishery Science Center, National Marine Fisheries Service, Honolulu, Hawaii |

**Table IV.**
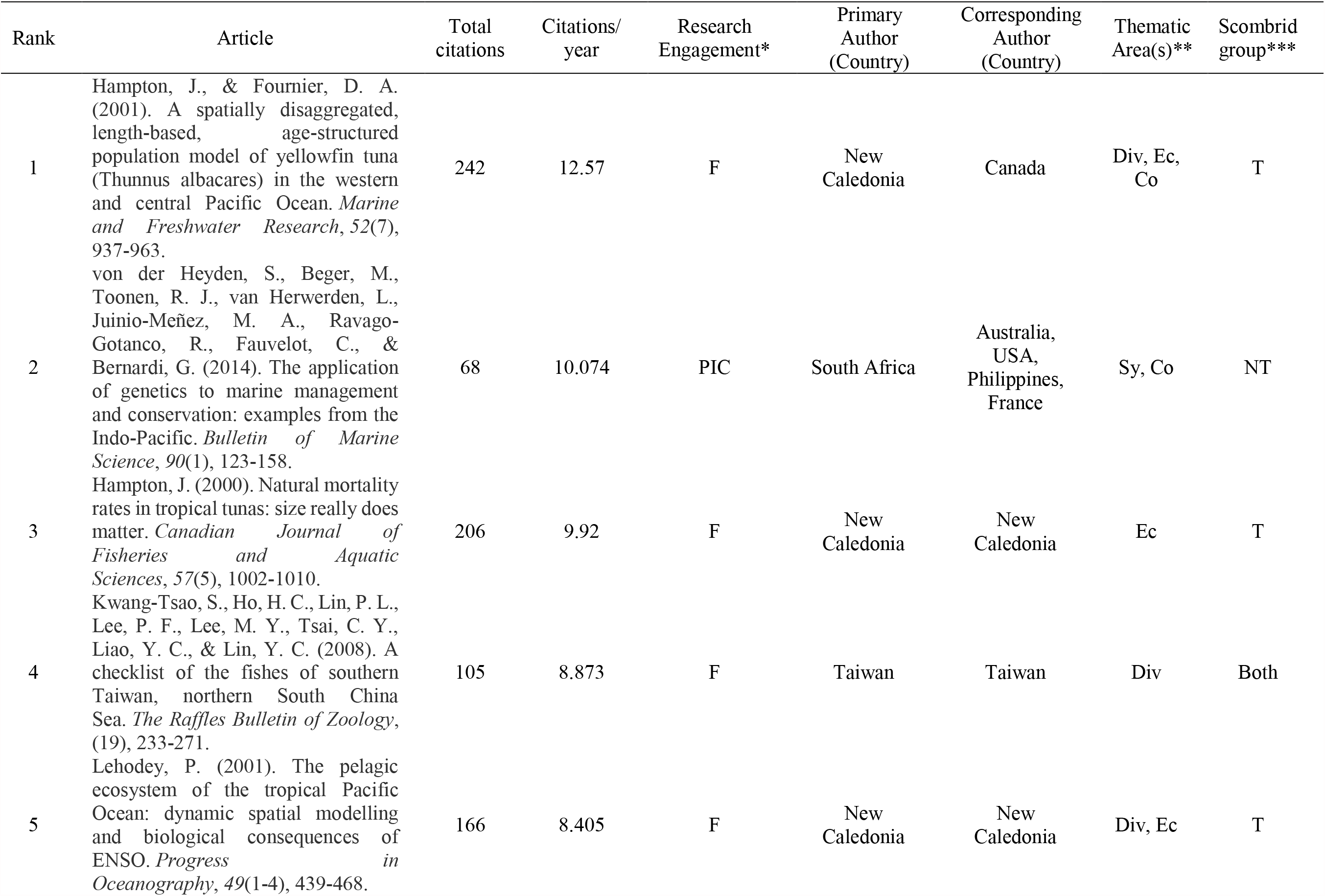
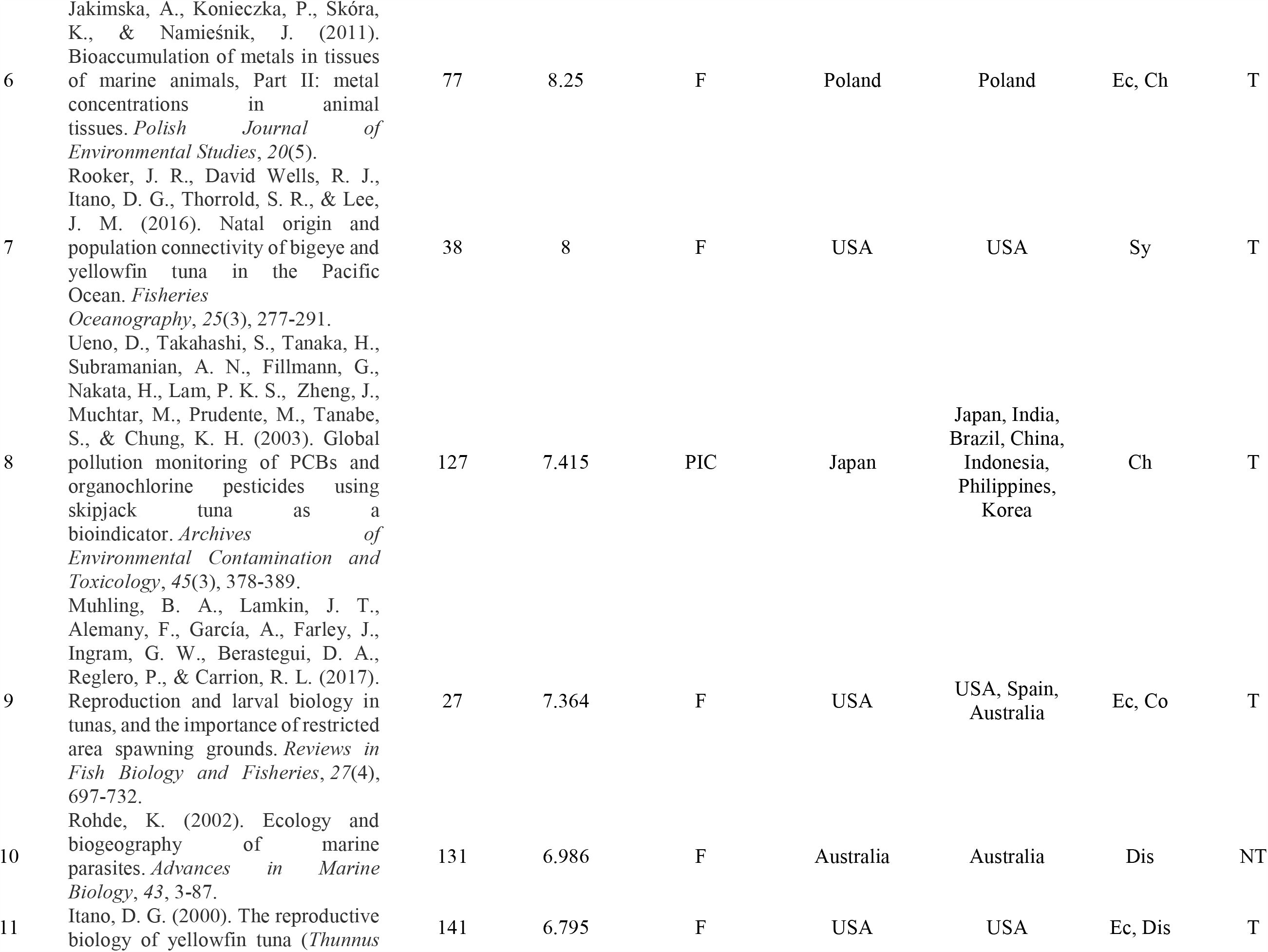
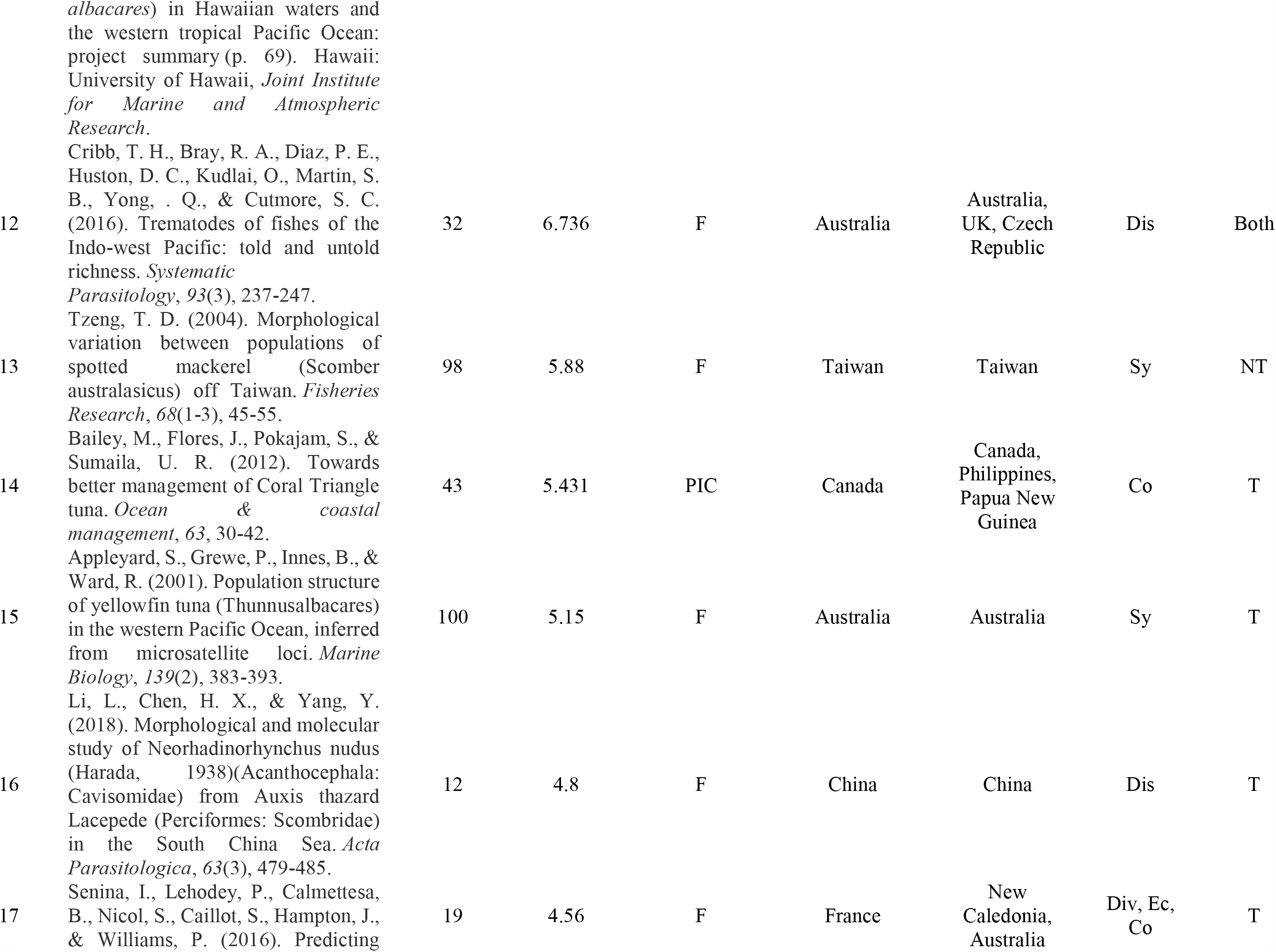
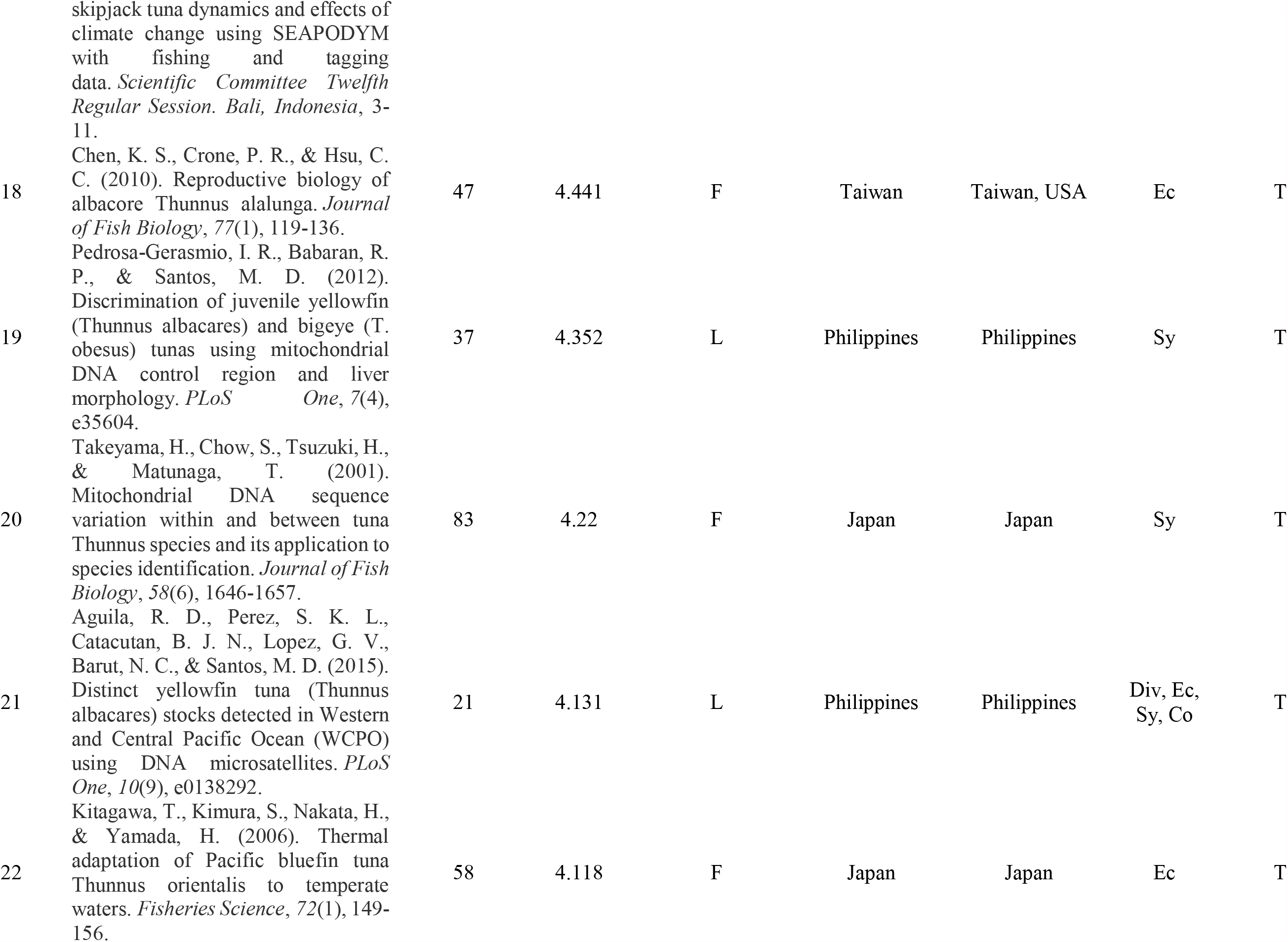
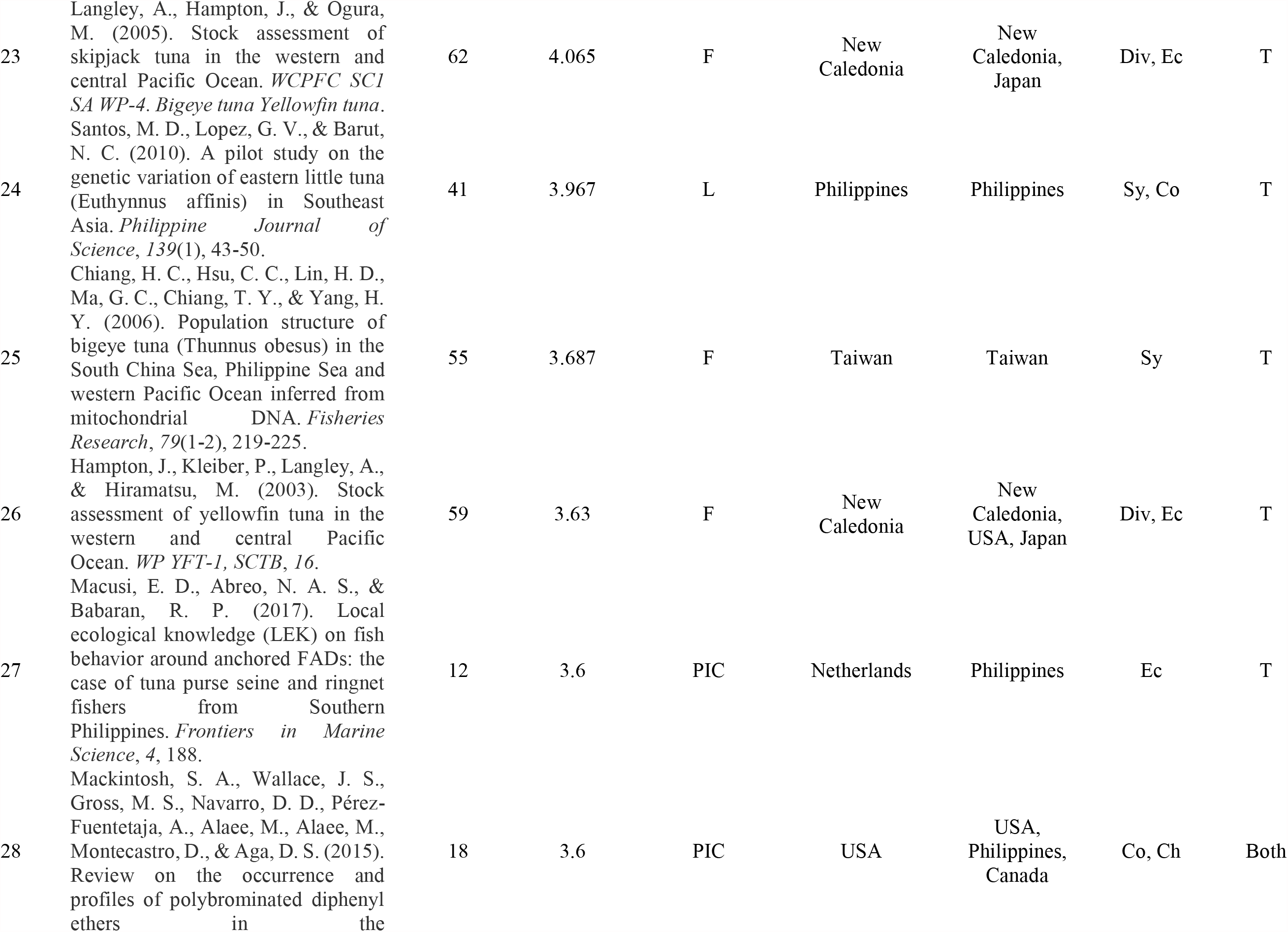
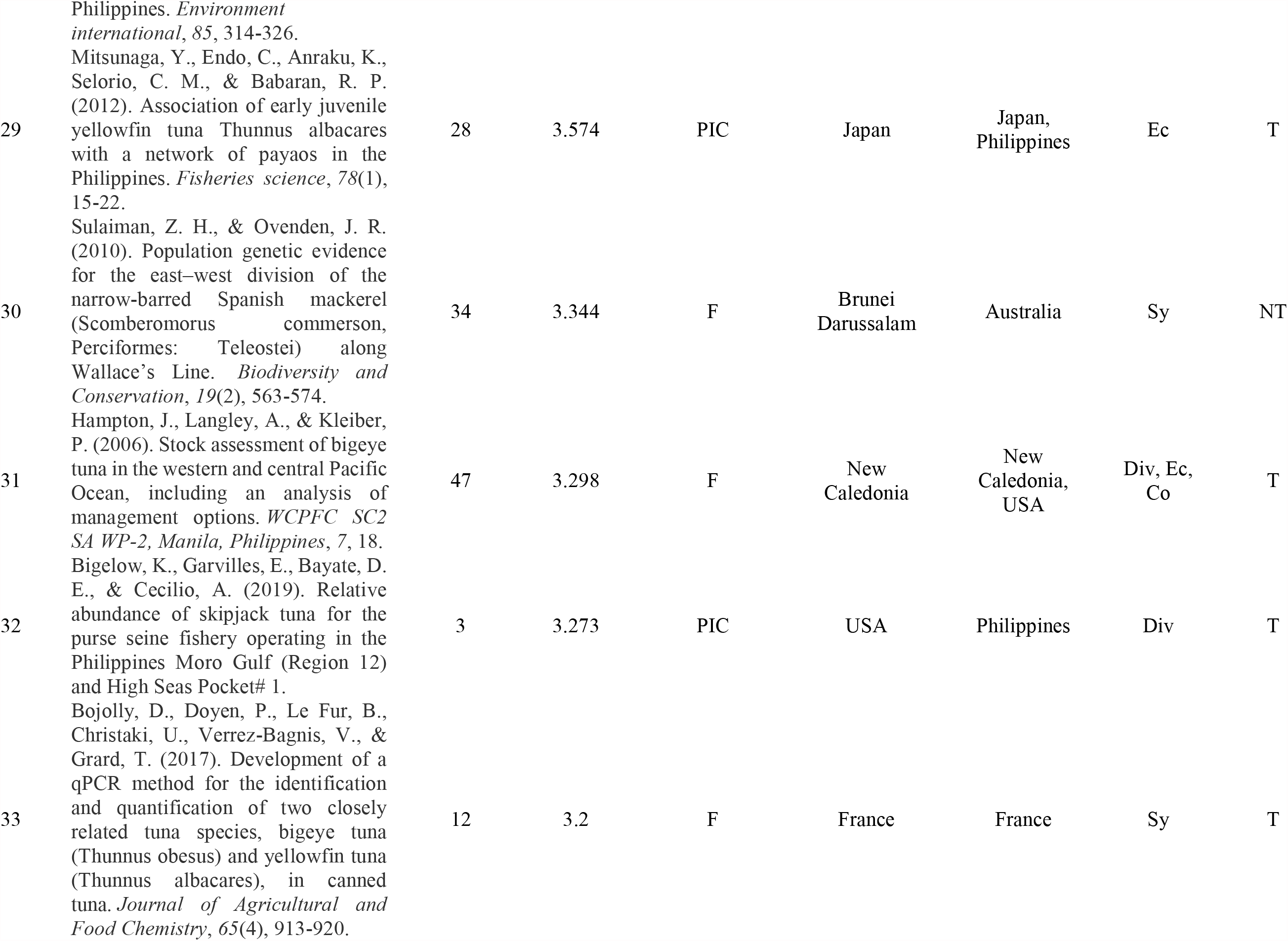
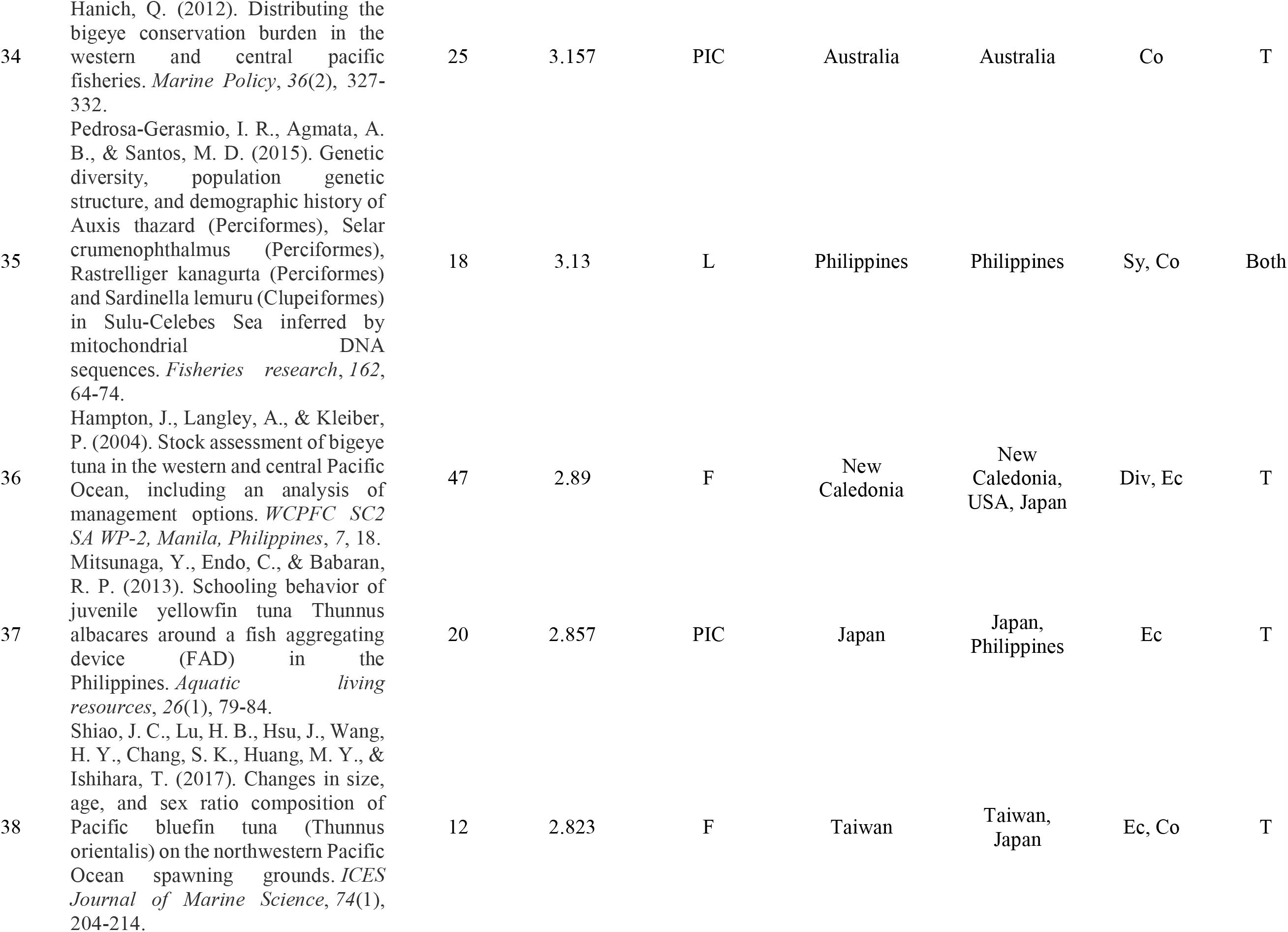
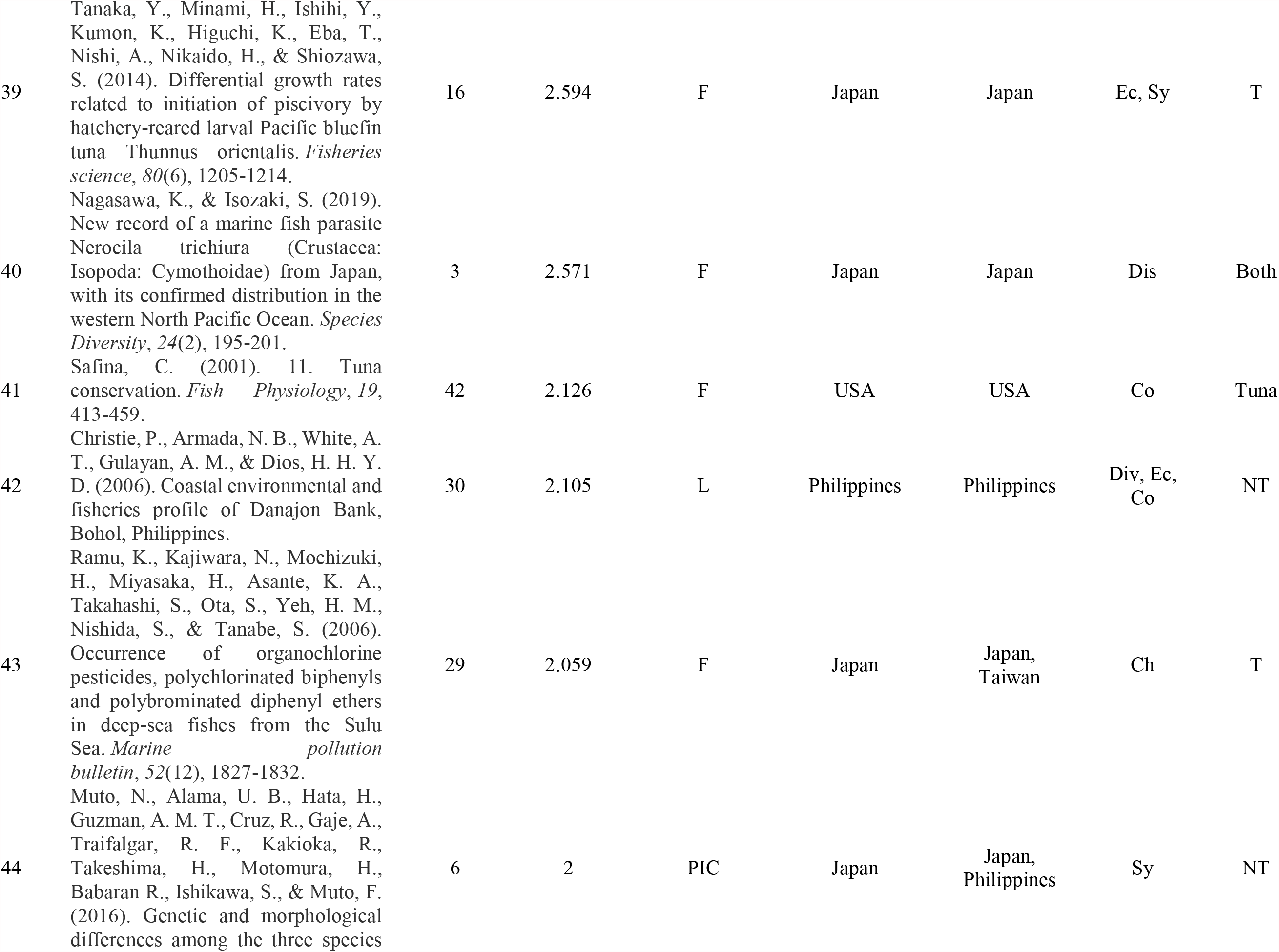
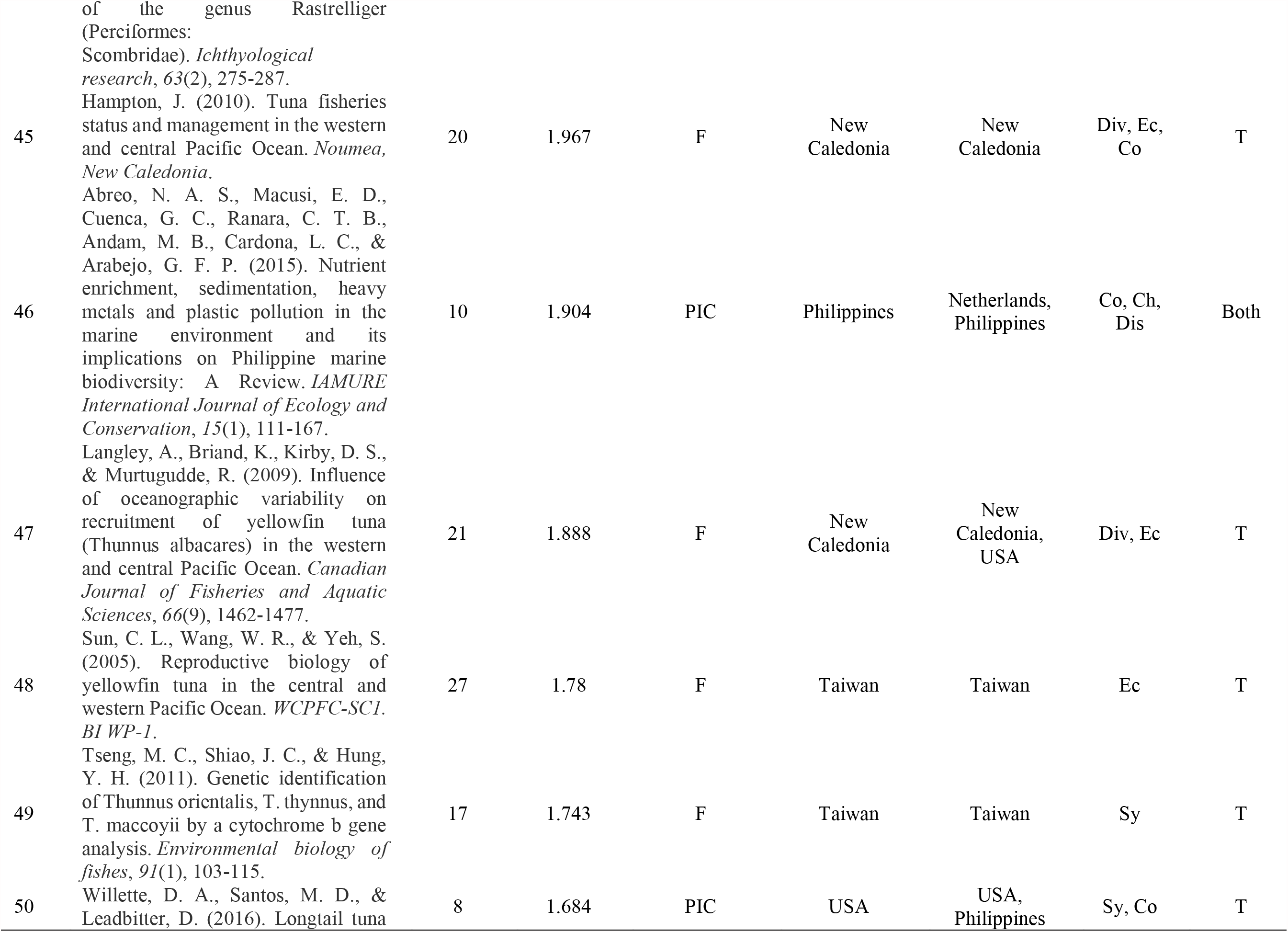
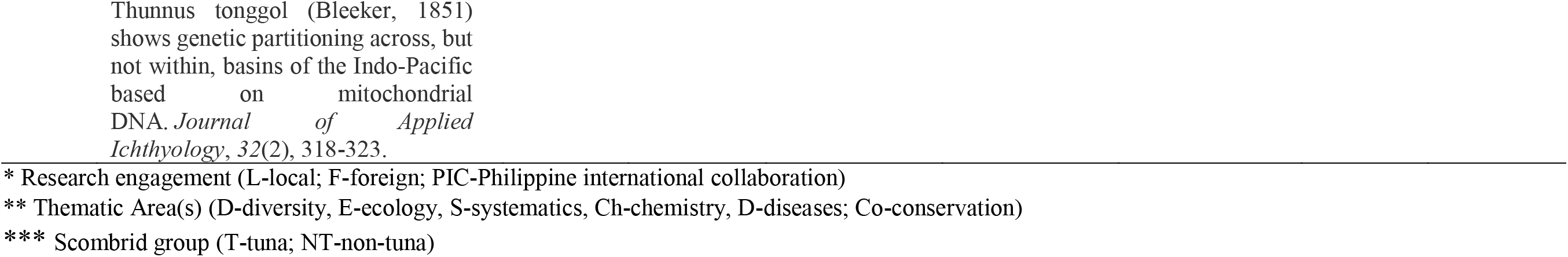
Top 50 most cited articles (Google Scholar) from 2000-2019.

## CONCLUSIONS

Information on Philippine scombrids based on the six thematic areas is crucial in updating both scientists and governments to address gaps in the R&D agenda, if necessary. As research outputs by Filipino researchers significantly lag their foreign counterparts in terms of citation index, international collaborations will undoubtedly increase impact and promote solidarity among nations towards its conservation and management. Should commercial exploitation persist without the commensurate research efforts to promote ecological management and conservation, a collapse of the Philippine scombrid population may not be too far from happening.

## Data Statement

Data used in the analysis are available upon request to the corresponding author.

## Author Contributions

J Gadut: data curation, methodology, resources, investigation, formal analysis, writing-original draft. C Deocaris: methodology, investigation, visualization, formal analysis, writing - review & editing. M Alinsug: conceptualization, resources, formal analysis, visualization, writing – original draft, writing - review & editing, supervision, funding acquisition.

## Acknowledgments

This study has been supported through a research grant from the Department of Science & Technology – Philippine Council for Agriculture, Aquatic & Natural Resources Research & Development (DOST-PCAARRD). The authors are grateful to the unwavering support of the various researchers & colleagues of MSU-GSC and staff of DOST-PCAARRD for the conduct of this research.

